# Dopamine promotes motor programs underlying substrate tunneling in larval *Drosophila*

**DOI:** 10.1101/2025.09.16.676552

**Authors:** James MacLeod, Bella Xu Ying, Maarten F. Zwart, Stefan R. Pulver

**Affiliations:** School of Psychology and Neuroscience, University of St Andrews, St Mary’s Quad, St Andrews, United Kingdom; Centre of Biophotonics, University of St Andrews, North Haugh, St Andrews, United Kingdom; Institute of Behavioural and Neural Sciences, University of St Andrews, St Andrews, United Kingdom

**Author notes:** Correspondence: Stefan R. Pulver. These authors contributed equally to this work (James MacLeod, Bella Xu Ying).

**Keywords:** *Drosophila melanogaster*, dopamine, locomotion, motor program, neuromodulation

## Abstract

Dopamine is a conserved biogenic amine with diverse neuromodulatory roles. Here we examine the role of dopamine in modulating larval *Drosophila melanogaster* motor programs underlying movement over and through substrates. First, we performed dual-color calcium imaging in tyrosine hydroxylase (TH)-expressing dopaminergic neurons and motor neurons to reveal cell-type specific recruitment patterns during fictive motor programs. Activity of select TH-neurons correlated strongly with fictive headsweeps as well as forward and backward locomotion. Next, bath applications of dopamine biased the isolated central nervous system towards fictive forward locomotion and inhibited fictive headsweeps. To probe whether these effects are recapitulated in intact animals, we optogenetically manipulated TH-neuron activity during surface crawling and tunneling. Optogenetic activation of TH-neurons with CsChrimson during crawling had no effect on headsweeps and slowed locomotor rhythms by increasing wave duration and decreasing wave frequency. Furthermore, posterior asymmetries, motor sequences characteristic of tunneling, were triggered. On the other hand, optogenetically inhibiting TH-neurons with GtACR1 had little effect on surface crawling. Underground, TH-neuron activation enhanced tunneling activity by increasing wave frequency, instead of duration, which increased overall time spent tunnelling, whereas inhibition decreased time spent tunneling. These results suggest that dopaminergic modulation of larval forward locomotion is dependent on sensorimotor context. We propose dopamine mediates a coordinated network effort to shift central pattern generators to promote tunneling-specific motor programs.

**New & Noteworthy:** Here, we find *Drosophila* larval dopaminergic neurons exhibit cell-type-specific activity in relation to fictive motor patterns. Furthermore, pharmacological and optogenetic manipulations of dopaminergic signaling acutely modifies locomotor output. Specifically, dopamine application promotes forward locomotion and inhibits headsweeps in isolated preparations, while optogenetically activating TH-neurons in freely behaving larvae mediates a shift from crawling to tunneling motor programs. This provides new insights into context-dependent dopaminergic modulation of locomotion in a genetically tractable invertebrate.

## Introduction

Dopamine is an evolutionarily ancient signaling molecule implicated in a wide range of processes and behaviors across phyla (1–3). Importantly, it has a prominent neuromodulatory role within motor systems. In both vertebrates and invertebrates, dopamine can activate multiple distinct receptor pathways and reconfigure motor networks to produce different motor programs (3–5). Despite decades of work in many model systems, our understanding of how dopamine reconfigures neural networks remains incomplete.

In vertebrates, dopamine influences many aspects of locomotion. Manipulation of dopamine levels often alters the rate of locomotion. For example, dopamine promotes locomotion in spinalized cats (6) and can induce fictive locomotion in the isolated rat spinal cord (7). Moreover, dopamine’s effects can be concentration dependent: in *Xenopus laevis* tadpole spinal preparations, application of low dopamine concentrations makes fictive locomotor episodes slower and more infrequent, while higher concentrations increase swim episode frequency (8). The current state of a locomotor network can also influence how dopamine affects locomotion. In the mouse spinal cord, the same concentration of dopamine can elicit different forms of fictive locomotion depending on the initial excitability state of the network (8). In invertebrates, such as the medicinal leech *Hirudo verbana*, dopamine switches fictive swimming rhythms into fictive crawling rhythms (9). Similarly, in the nematode worm *Caenorhabditis elegans*, dopamine is required for initiation and maintenance of gait transitions from swimming to crawling behaviors (9,10). Dopamine therefore appears to enable motor system transitions between activity states to facilitate adaptive behaviors across many species.

The *Drosophila melanogaster* larval locomotor system provides an attractive platform for understanding how dopamine can reconfigure locomotor networks. Approximately 120 dopaminergic neurons are present in the larval central nervous system (CNS) (11) and the distribution of dopamine receptors has been mapped (12). The roles that identified dopamine neurons play in larval learning and memory have also been extensively studied (13–16). However, less is known about the role of dopamine neurons in modulating larval motor programs. To date, dopamine’s overall effects on larval locomotor networks have only been partially characterized and yielded contradictory results. In some studies, dopamine appears to play a role in eliciting motor activity (17), in others, it suppresses locomotor activity or plays no discernible role in controlling movement (18,19). Previous work using calcium (Ca^2+^) imaging has demonstrated that signaling through muscarinic acetylcholine receptors (mAChRs) plays a critical role in generating rhythmic activity in thoracic and posterior regions of the larval CNS (20). This work suggests the presence of mAChR-dependent central pattern generators (CPGs) underlying forward/backward locomotion and headsweeps. To date, no studies have directly explored how dopamine modulates and reconfigures these CPG networks in the *Drosophila* larval CNS, nor how these effects translate to freely behaving larvae.

Here we combine dual-color Ca^2+^ imaging, pharmacology, and behavioral optogenetics to characterize how dopamine modulates locomotor output in *Drosophila* larvae. First, we perform dual-color imaging of cell activity in identified TH-expressing dopaminergic neurons and glutamatergic motor neurons in isolated larval CNS preparations to characterize dopaminergic activity dynamics during fictive motor behaviors. Next, we examine how pharmacological application of dopamine modulates locomotor CPG network activity in the isolated CNS. Finally, we optogenetically manipulate the activity of TH-neurons *in vivo* and examine the effects on crawling and tunneling. We find cell-type-specific recruitment during fictive locomotion and that dopamine differentially modulates motor program selection depending on the prevalent motor context. Specifically, activation of TH-neurons biases the motor network into tunneling by recapitulating tunneling-specific motor programs during crawling and heightening tunneling activity underground. Overall, this work characterizes dopamine’s role in promoting tunneling in *Drosophila* larvae and provides a foundation for understanding the diversity of dopamine’s actions within a genetically tractable locomotor network.

## Materials and Methods

### Animal rearing and genetic constructs

*Drosophila* were raised on standard cornmeal-based fly food and incubated at 20°C with ∼55-60% humidity. Feeding third-instar larvae were used in all experiments. For dual-color experiments, the green-shifted Ca^2+^ indicator GCaMP6f (21) was expressed in dopaminergic neurons in the TH-GAL4 expression pattern (22). The red-shifted Ca^2+^ indicator jRCaMP1b (23) was expressed in glutamatergic neurons (which includes all motor neurons) in the vGlut-LexA expression pattern (24). For pharmacological experiments, GCaMP3 was expressed in the OK371-GAL4 expression pattern (25) to label motor neurons. Finally, for optogenetic experiments, to activate TH-neurons, the excitatory red-light gated cation channel CsChrimson was expressed in dopaminergic neurons in the TH-GAL4 expression pattern (*UAS-20X-CsChrimson-mVenus* in the attP2 insertion site). To inhibit TH-neurons, red-light gated anion channel GtACR1 was expressed (*UAS-20X-GtACR1-mVenus* in the VK00005 attP insertion site). Optogenetic experimental animals were raised on media supplemented with 0.5mM all trans-retinal (Sigma-Aldrich, Dorset, UK) and kept in darkness, whereas control animals were raised on media without retinal.

### Dual-color Ca^2+^ imaging of TH-neurons and motor neurons

Larvae (Fig. 1A) were placed in Sylgard-lined dissecting dishes with physiological saline containing (in mM) 135 NaCl, 5 KCl, 4 MgCl_2_·6H_2_O, 2 CaCl_2_·2H_2_O, 5 TES buffer, 36 sucrose, pH 7.15. Animals were pinned to the Sylgard dish with 0.1mm tungsten wire (California Fine Wire, CA, USA) through the head and posterior segment. Two transverse incisions were made near the rostral and caudal pins with microdissection scissors. Then, a longitudinal cut connecting the transverse cuts was made, and the body wall was pinned down with 0.01mm tungsten wire (Fig. 1B). Internal organs and trachea were removed with forceps. The CNS was dissected away from the body wall and placed on Sylgard-lined dishes secured with fine tungsten wire while submerged in physiological saline.

**Figure 1.**
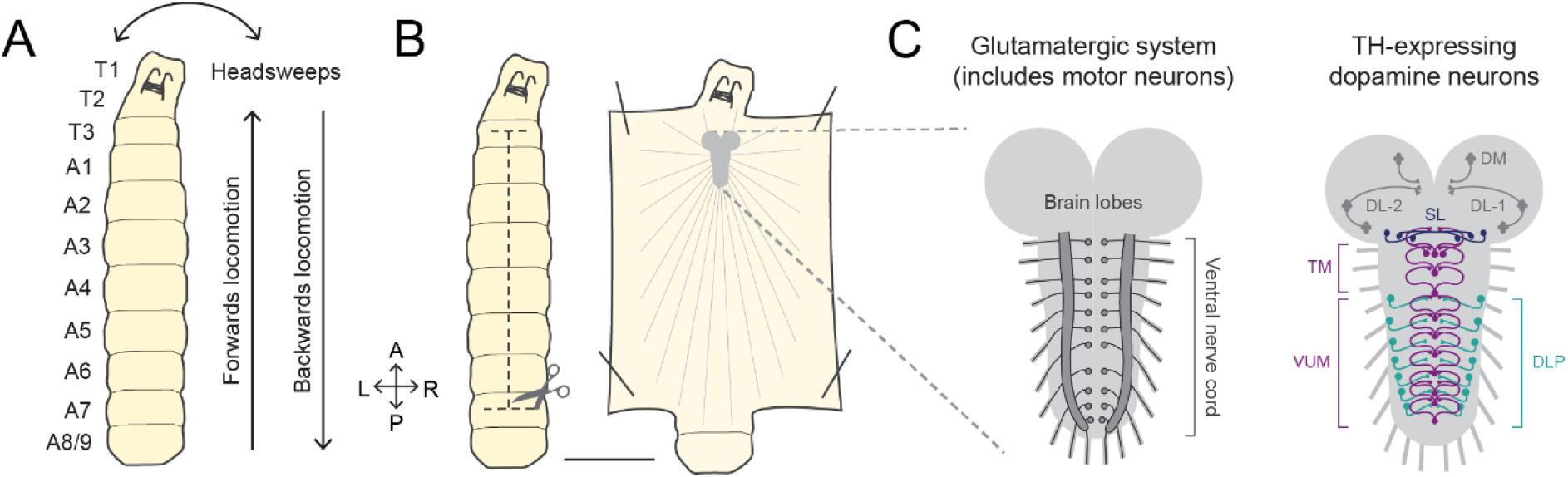
*Drosophila* larval anatomy, motor programs and central nervous system. **A:** Diagram of third-instar *Drosophila* larva. T1-T3 are thoracic segments, A1-A8/A9 are abdominal segments. Common larval motor programs are labelled. **B:** Dissection method for isolated CNS Ca^2+^ imaging experiments. One longitudinal and two transverse cuts (dashed lines) are made before the body wall is pinned down and CNS is removed. Scale bar = 1mm. **C:** Isolated CNS (brain lobes and ventral nerve cord (VNC)) showing glutamatergic system on the left (circles: motor neuron somas, filled lateral areas: neuropils, lines: axons) and TH-expressing neurons on the right (DM and DL clusters in the brain lobes (grey), SL neurons in the suboesophageal zone (dark blue), TM and VUM neurons in the medial VNC (purple) and DLP neurons in the lateral VNC (cyan)). DM: dorsomedial, DL: dorsolateral, SL: suboesophageal, TM: thoracic medial, VUM: ventral unpaired medial, DLP: dorsolateral paired.

Motor neurons and TH-neurons (Fig. 1C) were imaged using a custom-built spinning disk confocal microscope with a XLUMPLFLN 20X dipping objective (Olympus, Tokyo, Japan), SOLA light engine (Lumencor, OR, USA) and an ORCA-Fusion CMOS camera (Hamamatsu, Hamamatsu, Japan) (Fig. 1D). Light emitted from the fluorophores was split by wavelength using dichroic mirrors in a Cairn OptoSplit beam-splitter (Cairn Instruments, Kent, UK). Preparations were imaged using an X-light spinning disk confocal with a 70µm pinhole array (Crest Optics, Rome, Italy) for 5-10min at 5fps with 150ms exposure time. Preparations with ambiguous cell-type identity were further imaged in wide-field for 1-2min with a manual z-axis scan to aid identification.

Fluorescence intensity values were extracted from motor neurons and select TH-neurons from regions of interest (ROIs) drawn in Fiji (26). Custom Python scripts were used to perform correlations and waveform analysis to identify forward/backward waves and headsweep events. Fictive forward locomotion was defined as waves of activity beginning in A8 and propagating to T2-3. Fictive backward locomotion was defined as waves beginning in T2-3 and progressing to A8. Fictive headsweeps were defined as bilaterally asymmetric activity in thoracic regions: ROIs were drawn on the three thoracic hemi-segments on both hemispheres (6 total) to characterize bilaterally asymmetric activity. Average intensity values in each right thoracic ROI were then subtracted from the left. Fictive headsweeps were defined as peaks greater than ±1.8SD from the mean ΔF/F. Pearson’s correlations between dopamine and fictive locomotion traces were performed in 80-200s periods. Time-lagged cross-correlation consisted of series of correlation calculations carried out with one trace shifted at 200ms intervals relative to another over a 10s window. Correlation and time lag boxplots were plotted in Origin Pro.

### Pharmacological applications of dopamine and L-dopa

Isolated CNS preparations were performed as above. For dopamine bath applications, different concentrations (100µM, 250µM and 500µM) of dopamine hydrochloride (Sigma-Aldrich, Dorset, UK) were dissolved in physiological saline and superfused at 1-2ml/min. Control periods of 6-10min in saline were imaged, then dopamine was superfused over the isolated CNS for 6-10min, with a final wash period of 6-10min in saline. The final 4min of each condition were analyzed. For L-dopa, semi-intact preparations were first preincubated for 20min in either: 500µM L-dopa (Sigma-Aldrich, Dorset, UK) dissolved in physiological saline with equimolar ascorbic acid, or saline (control). Then, the CNS was isolated and imaged for 10min; the full imaging period was analyzed.

A customized BX51WI fluorescent compound microscope (Olympus Microscopes, Tokyo, Japan) mounted on an X-Y translation stage (Siskiyou Corporation, OR, USA). A dual-channel LED controller (Cairn Research Ltd., Kent, UK) was used as a light source for both transmitted and fluorescence light. Images were acquired using an R1 Retiga camera (Qimaging, Surrey, Canada) and μManager (27). Mean ROI fluorescence values were obtained in Fiji. Waveform analysis was performed with custom Python scripts to identify different fictive events. Posterior bursts were defined as waves of activity only propagating from A8 to A5. Other fictive behaviors during dopamine application were defined as in *Dual-color Ca^2+^ imaging of <u>TH-neurons and motor neurons</u>*. For L-dopa experiments, the headsweep threshold was based on absolute differences in ΔF/F. Unilateral thoracic activity peaks not matched by peaks on the contralateral side were detected as headsweeps. To account for negative values, the trace was rectified and the area under the curve was measured. For dopamine, statistical analyses were carried out as Mauchly’s tests of sphericity and one-way repeated-measures ANOVAs with Tukey HSD pairwise comparisons. For L-dopa, Kolmogorov-Smirnov normality tests and Welch’s t-tests were used due to different sample sizes (control n = 11 vs L-dopa n = 8). All statistics and scatter/boxplots were plotted in Origin Pro.

### Optogenetics on freely behaving larvae

All experiments were performed in darkness. Behavioral arenas were illuminated from below with 850nm infrared light (Andoer IR49S Mini). Optogenetic stimulation was delivered via a 998mA Thorlabs High-power Solis white LED controlled with a DC2200 LED Driver (Thorlabs Inc, NJ, USA). Off-on-off periods for stimulation light lasted 30s-40s-30s for all experiments. For surface crawling experiments, larvae were placed on a 13cm circular dish filled with 1% (w/v) agarose (Fisher Scientific, Loughborough, UK) (Fig. 4A). Larvae were allowed to acclimate for 2min before recording. Videos were recorded at 15fps with a Ximea MQ013RG-E2 camera (Ximea GmbH, Münster, Germany) attached to a Navitar ZOOM 7000 macro lens (Navitar Inc, NY, USA). For tunneling experiments, 5cm circular dishes were filled with 1% agarose and a small divot was carved at a 30° angle (Fig. 5A). Larvae were brushed into the divot headfirst and allowed to acclimate for 1min before recording. Larvae that tunneled to the bottom of the dish were discarded. Experiments were repeated for (crawling n = 8 and tunneling n = 10) larvae per genotype.

Crawling videos were background-subtracted with a custom Fiji macro before behavioral tracking with FIMTrack (28). Wave frequency was defined as number of waves divided by total time. Mean wave duration was calculated as total crawling time (total time – (headsweeping time + time at 0mm/s velocity)) divided by wave frequency. Left headsweeps were defined as a ≥210° body angle and right headsweeps a ≤150° body angle, with straight larvae being 180°. Headsweep duration represented total time spent matching above criteria. Tunneling videos were downsampled to every second frame and waves were manually tracked. Posterior asymmetries were also manually annotated for both crawling and tunneling videos. In crawling, posterior asymmetries were defined as posterior segment bends of >10° (<u>Supplementary Video</u> <u>S1</u>). In tunneling, they represented posterior segment bends in a different orientation to the main body length (<u>Supplementary Video S2</u>) due to less space underground.

Statistical analysis was conducted as Mauchly’s tests of sphericity, Shapiro-Wilk normality tests and Levene’s tests of homoscedasticity. Since no behavioral parameters met all three assumptions, Greenhouse-Geisser corrections were applied to mixed-factorial ANOVAs and Bonferroni corrections to pairwise comparisons. Behavioral analyses and statistics were performed in Python. Box plots were plotted in Origin Pro.

## Results

### TH-neurons are differentially recruited into locomotor activity

To determine how activity in TH-neurons is linked to locomotor activity, we simultaneously imaged Ca^2+^ dynamics in glutamatergic motor neurons and TH-neurons in isolated CNS preparations during fictive locomotion. The TH-GAL4 driver line labels distinct, identifiable dopamine neuron populations in the brain and VNC (Fig. 1C). Bilaterally symmetric DL1, DL2 and DM clusters are found in the brain lobes, and the SL neurons are found in the suboesophageal zone (SEZ) (11). Based on morphology and location there are two subtypes of dopaminergic neuron in the VNC. Abdominal segments A1-7 contain two DLP neurons and one VUM neuron. VUM neurons in thoracic segments are referred to as TM neurons (29). Thoracic segments contain one TM neuron each and no DLP neurons, except for segment T1 which contains three TM neurons.

SL neurons project both ipsilaterally and contralaterally (30). Comparing activity in the processes and somas showed that the contralateral projections were visible, whereas ipsilateral processes projected ventrally and were not visible. The SL neuron pair showed clear periods of asymmetric activity where one SL neuron was more active than the other. Therefore, we compared asymmetric activity in SL neurons with glutamatergic asymmetric activity in thoracic segments, a measure of fictive headsweeps. SL bilateral asymmetry did not correlate strongly with glutamatergic asymmetries (Pearson’s r range = -0.26 to 0.20, n = 3) (Fig. 2Ai). Then, we compared SL asymmetries to the rate of change of glutamatergic asymmetry during fictive headsweeps (i.e., the change in ΔF/F difference between left and right thoracic segments per second) (Fig. 2Aii). Asymmetric SL activity was positively correlated to the rate of change of asymmetric glutamatergic activity (mean Pearson’s r = 0.68, n = 3) (Fig. 2Aiii). Time-lagged cross-correlation between SL asymmetry and the rate of change of glutamatergic asymmetry showed no consistent time lag (mean = -0.07s), suggesting SL activity and the rate of change of glutamatergic activity occur nearly simultaneously (Fig. 2Aiii).

**Figure 2.**
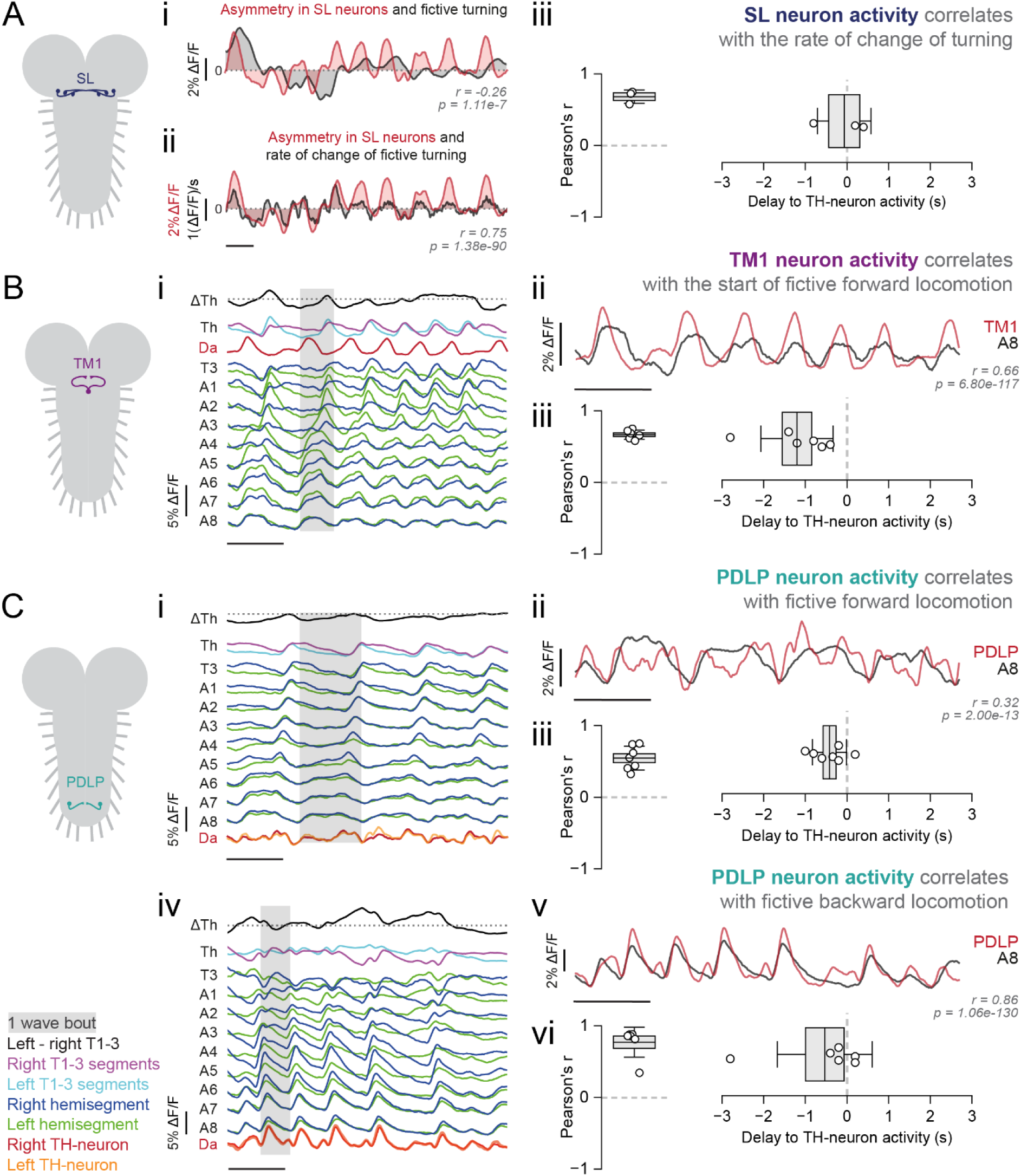
TH-neurons are differentially recruited into locomotor activity. Ai: Asymmetry in SL neuron activity (red) relative to asymmetry in thoracic motor neuron activity (black) during fictive headsweeps (r = -0.26, p = 1.11e-7). Dotted line at 0 = symmetrical activity. **Aii:** Asymmetry in SL neuron processes (red) relative to rate of change of thoracic motor neuron asymmetric activity (black) during fictive headsweeps (r = 0.75, p = 1.38e-90). **Aiii:** Pearson’s correlation and delay of highest correlation in time-lagged cross-correlation between SL neuron asymmetry and rate of change of glutamatergic thoracic asymmetric activity; n = 3. **Bi:** ΔF/F trace of TM1 neuron activity (red) relative to segmental motor neuron activity. Th = mean activity across thoracic segments, ΔTh = difference between left/right thoracic ΔF/F to illustrate asymmetric activity, T3-A8 = raw ΔF/F. **Bii:** ΔF/F trace of TM1 neuron activity (red) correlated to A8 motor neuron activity (black) during fictive forward locomotion (r = 0.66, p = 6.80e-117). **Biii:** Pearson’s correlation and time-lagged cross-correlation between TM1 neuron activity and A8 activity during fictive forward locomotion, n = 6. **Ci and iv:** ΔF/F trace of PDLP neuron activity (red) relative to segmental motor neuron activity. **Cii and v:** ΔF/F trace of PDLP neuron activity (red) correlated to A8 motor neuron activity (black) during fictive forward locomotion (r = 0.32, p = 2.00e-13) and fictive backward locomotion (r = 0.86, p = 1.06e-130). **Ciii and vi:** Pearson’s correlation and time-lagged cross-correlation between PDLP neuron activity and A8 activity during fictive forward (n = 7) and backward locomotion (n = 6). Boxplots = mean ±1SEM., whiskers = ±1SD. All time scale bars = 20s.

The central TM neuron in segment T1 (TM1) is a large neuron with extensive arborizations projecting to the medial and lateral thoracic regions. During fictive forward locomotion, TM1 neurons were active immediately before glutamatergic activity in segment A8, i.e., at the beginning of each forward wave (Fig. 2Bi and ii). TM1 activity correlated strongly with A8 activity (mean Pearson’s r = 0.67, n = 6) and activity increased prior to the first forward wave of a bout in 100% of instances by an average of -1.2s (Fig. 2Biii).

The most posterior DLP neurons (PDLP) in segment A7 were easily identifiable, and their location as the rearmost dopamine neurons makes them candidates for involvement in forward wave initiation (Fig. 2Ci-ii and iv-v). PDLP neurons showed positive correlations with A8 glutamatergic activity during both fictive forward (mean Pearson’s r = 0.54, n = 7) (Fig. 2Ciii) and fictive backward locomotion (mean Pearson’s r = 0.77, n = 6) (Fig. 2Cvi). Time-lagged cross-correlation suggested that PDLP activity may precede local glutamatergic activity during both fictive forward (mean time lag = -0.42s) (Fig. 2Ciii) and backward locomotion (mean time lag = -0.53s) (Fig. 2Cvi).

### Dopamine promotes fictive forward locomotion and suppresses headsweeps in a concentration-dependent manner

Given the clear relationships between TH-neuron activity and fictive motor programs, we wanted to examine how dopamine directly modulates CPG networks underlying locomotion in *Drosophila* larvae. We therefore measured CPG activity in the isolated larval CNS in the presence of dopamine (Fig. 3A and B). Application of 250µM and 500µM dopamine significantly increased the number of fictive forward locomotor waves (Control-250µM: q(12) = 9.1, p<1e-4; 250µM-Wash: q(12) = 9.6, p<1e-4; Control-500µM: q(16) = 9.9, p<1e-4; 500µM-Wash: q(16) = 9.2, p<1e-4) (Fig. 3C), while only 250µM significantly decreased fictive backward wave frequency (Control-250µM: q(12) = 4.5, p = 0.019; 250µM-Wash: q(12) = 5.7, p = 4.6e-3) (Fig. 3D). Similarly, the two highest concentrations severely dampened the frequency of fictive headsweeps (Control-250µM: q(12) = 9.1, p<1e-4; 250µM-Wash: q(12) = 9.6, p<1e-4; Control-500µM: q(16) = 9.9, p<1e-4; 500µM-Wash: q(16) = 9.2, p<1e-4) (Fig. 3C). Application of 100µM dopamine did not significantly alter any motor program frequency, and dopamine had no effects on posterior burst frequency. Since endogenous dopamine in the CNS may not change uniformly during locomotion, we also used L-dopa to manipulate global dopamine in a more physiologically relevant manner (Supplementary Fig. S1A). As with 250µM and 500µM dopamine, 500µM L-dopa increased fictive forward wave frequency (t(7.6) = -4.7, p = 1.8e-3) (Supplementary Fig. S1B) and suppressed headsweeps (t(11.6) = -5.1, p = 3.2e-4) (Supplementary Fig. S1C).

**Figure 3.**
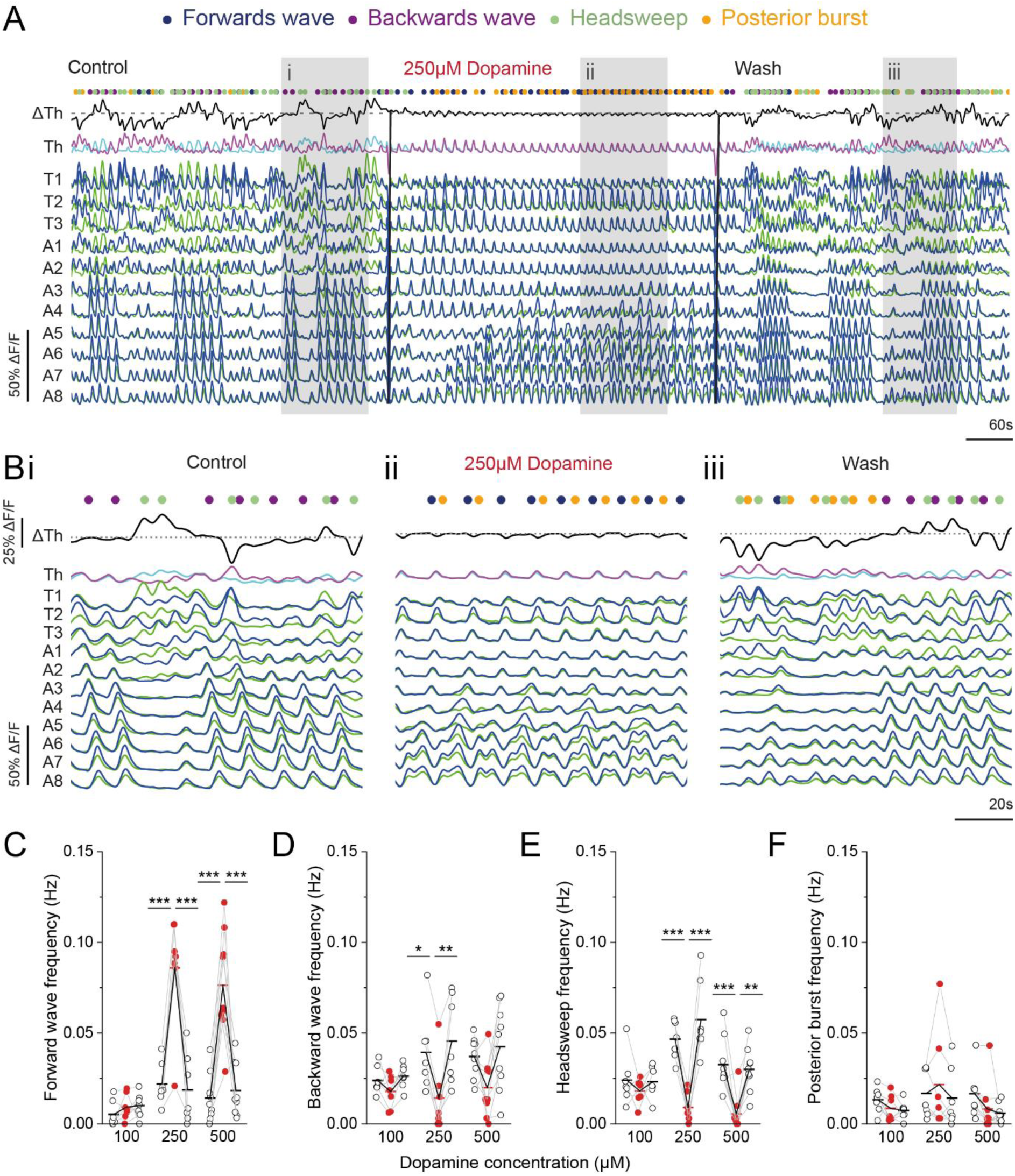
Higher dopamine concentrations promote fictive forward locomotion and suppress headsweeps. **A:** Full ΔF/F trace of segmental motor neuron activity during control saline, 250µM dopamine and wash saline applications. Th = mean activity across thoracic segments, ΔTh = difference between left/right thoracic ΔF/F to illustrate asymmetric activity, T3-A8 = raw ΔF/F. Dots = detected fictive motor program events. **B:** Expanded views of grey sections in A. **C-F:** Scatter plots comparing motor program frequency across dopamine concentrations. **C:** fictive forward waves, **D:** fictive backward waves, **E:** headsweeps, **F:** posterior bursts. Thick horizontal black/red lines = mean. One-way repeated-measures ANOVA, ***p<0.001, **p<0.01, *p<0.05. 100µM: n = 7, 250µM: n = 7, 500µM: n = 9.

**Supplementary Figure S1.**
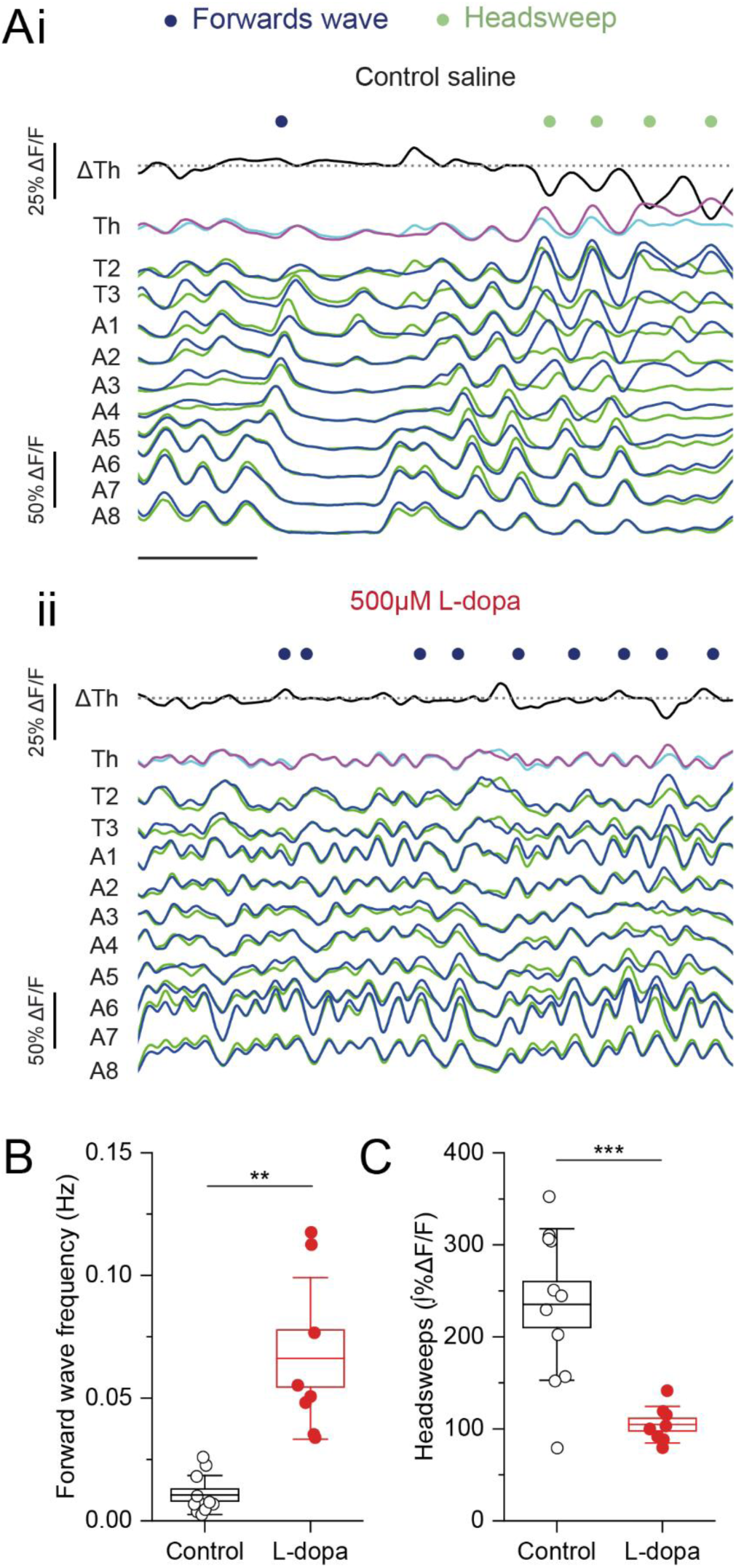
Preincubation in L-Dopa promotes fictive forward locomotion and suppresses headsweeps. **A:** ΔF/F trace of segmental motor neuron activity in i: control saline and ii: 500µM L-dopa. Th = mean activity across thoracic segments, ΔTh = difference between left/right thoracic ΔF/F to illustrate asymmetric activity, T3-A8 = raw ΔF/F. Dots = detected fictive motor program events. Scale = 20s. **B and C:** Fictive forward wave frequency and headsweeps in control saline vs 500µM L-dopa. Welch’s t-test, ***p<0.001, **p<0.01. Control: n = 11, L-dopa: n = 8. Boxplots = mean ±1SEM., whiskers = ±1SD.

### TH-neurons promote tunneling behaviors

After examining dopamine’s effects on the isolated CNS during fictive locomotion, we wanted to understand if these effects are recapitulated in freely behaving animals. To this end, we optogenetically activated or inhibited all dopamine neurons in the TH-GAL4 expression pattern with CsChrimson or GtACR1, respectively, during surface crawling (Fig. 4B). Activation of TH-neurons increased mean forward wave duration (Before-During: T(7) = -3.8, p = 0.02; During-After: T(7) = -3.16, p = 0.048; During(-retinal)-During(+retinal): T(14) = -3.9, p = 0.031) (Fig. 4C) and decreased forward wave frequency (Before-During: T(7) = 6.6, p = 9e-4; During-After: T(7) = 4.4, p = 9.9e-3; During(-retinal)-During(+retinal): T(14) = 5.6, p = 1.3e-3) (Fig. 4D). Meanwhile, inhibiting TH-neurons had no effects on wave duration nor frequency (Fig. 4C and D). Activating and inhibiting TH-neurons both had no effect on headsweep frequency (Fig. 4E). Backward wave frequency also did not differ between control and experimental animals. In addition, we identified and characterized a novel form of behavior, not previously documented in crawling/tunneling assays, termed “posterior asymmetries”, defined as a spontaneous posterior lateral bend away from straight body angle (Fig. 4F) (<u>Supplementary Video S1</u>). These events rarely occurred in no retinal controls and in light-off periods but significantly increased in experimental larvae during TH-neuron activation (Before-During: T(7) = -7.7, p = 3.6e-4; During-After: T(7) = -5.5, p = 3e-3) (Fig. 4G).

**Figure 4.**
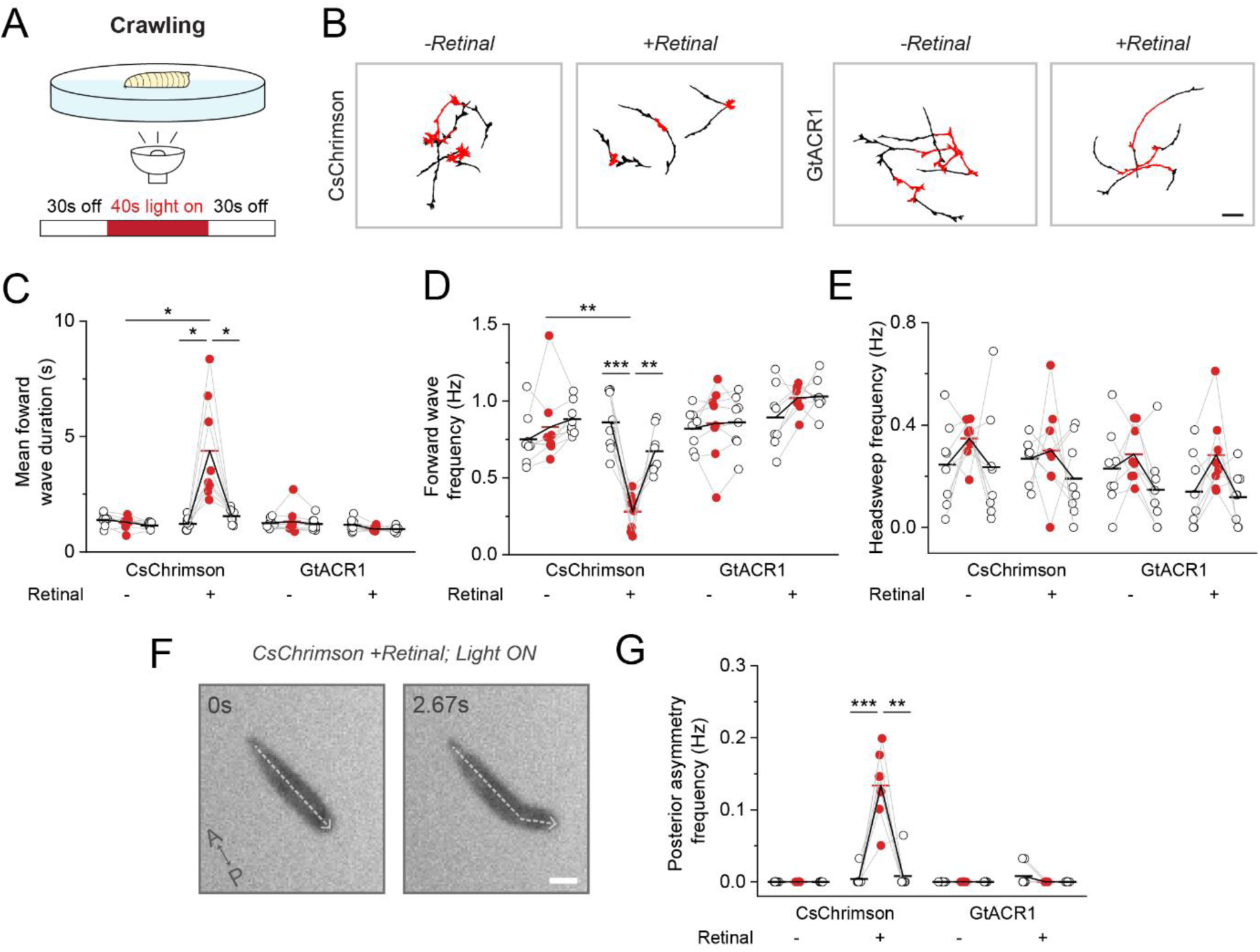
Activating TH-neurons during crawling slows forward locomotor rhythms and promotes posterior asymmetries. **A:** Diagram of optogenetic protocol during crawling. **B:** Walk paths of control and experimental animals for CsChrimson (activation) and GtACR1 (inhibition) during light-off (black) and light-on (red) periods. Scale bar = 1cm. **C-E and G:** Scatter plots comparing locomotor parameters across conditions. **C:** mean wave duration, **D:** wave frequency, **E:** headsweep frequency and **G:** posterior asymmetry frequency. Thick horizontal black/red lines = mean. Mixed-factorial ANOVA with Bonferroni-corrected post hoc analysis, ***p<0.001, **p<0.01, *p<0.05, n = 8. **F:** Image sequences showing experimental larvae performing posterior asymmetry upon TH-neuron activation. Scale bar = 1mm.

The effects seen when activating TH-neurons during crawling (slower wave duration, lower wave frequency) were reminiscent of features observed during larval tunneling (31). Nevertheless, 0% of larvae attempted to tunnel upon optogenetic activation during crawling (n = 8). To examine potential differences in dopaminergic modulation when larvae are tunneling (different motor context), we performed the same optogenetic manipulations with larvae tunneling in agarose (Fig. 5B). Interestingly, forward wave duration remained unchanged across stimulation conditions during tunneling (Fig. 5C). However, activating TH-neurons significantly increased forward wave frequency compared to both controls and the pre-light period (Before-During: T(9) = -3.2, p = 0.03; During(-retinal)-During(+retinal): T(18) = -5.2, p = 1e-3) (Fig. 5D), whereas inhibiting TH-neurons had no effect. Backward wave frequency did not differ between control and experimental animals. Activating TH-neurons also increased the total time spent tunneling, compared to controls and the pre-light period (Before-During: T(9) = -3.9, p = 0.011; During(-retinal)- During(+retinal): T(18) = -6.0, p = 2e-4) (Fig. 5E). On the other hand, inhibiting TH-neurons significantly decreased time spent tunneling compared to the pre-light period (Before-During: T(9) = 6.2, p = 4.9e-4) (Fig. 5E). Finally, although posterior asymmetries are normally almost exclusively found during tunneling (<u>Supplementary</u> <u>Video S2</u>), optogenetic activation/inhibition had no effect on posterior asymmetry frequency (Fig. 5F).

**Figure 5.**
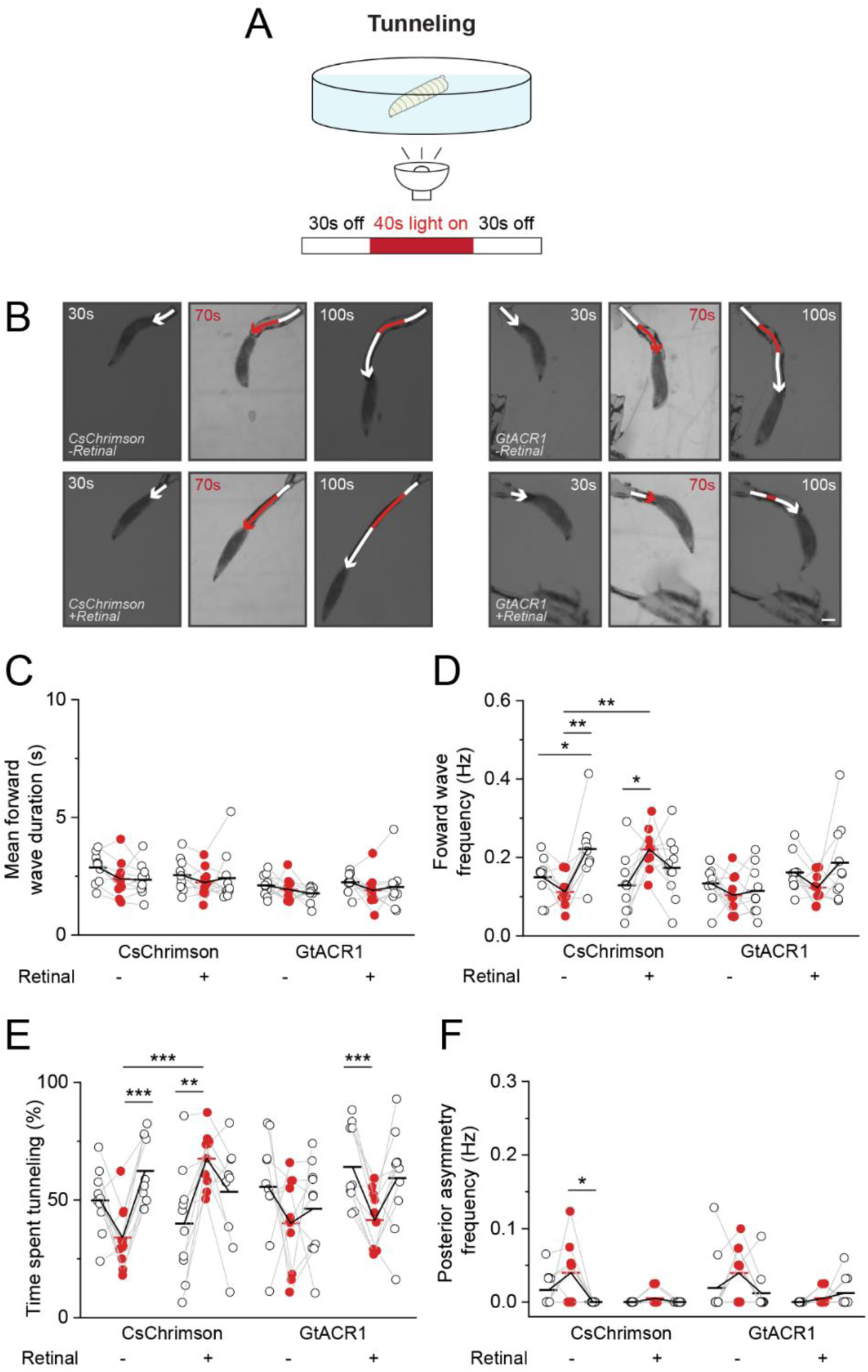
Activating TH-neurons underground promotes tunneling. **A:** Diagram of optogenetic protocol during tunneling. **B:** Images at end of before-during-after periods showing tunneling progress; light-off (white arrows) and light-on (red arrows). Scale bar = 1mm. **C-F:** Scatter plots comparing locomotor parameters across conditions. **C:** mean wave duration, **D:** wave frequency, **E:** total time spent tunneling and **F:** posterior asymmetry frequency. Thick horizontal black/red lines = mean. Mixed-factorial ANOVA with Bonferroni-corrected post hoc analysis, ***p<0.001, **p<0.01, *p<0.05, n = 10.

## Discussion

Here, we combined various methods to examine how dopamine modulates locomotor CPG networks in *Drosophila* larvae. We studied dopamine modulation across multiple levels of analysis, from imaging TH-neurons during fictive motor activity to pharmacological manipulations in isolated CNS preparations and optogenetics in intact, freely behaving animals. Using complementary approaches, we have shown how dopamine shapes fictive and real locomotion. First, we uncovered surprising diversity in the recruitment patterns of TH-neurons. Motor programs for forward/backward locomotion and headsweeps preferentially recruited specific TH-neurons, whose activity encoded localized motor activity. Moreover, across isolated CNS experiments, high concentrations of dopamine consistently suppressed the generation of bilaterally asymmetric activity associated with the headsweep motor program. This supports the idea that dopamine plays a key role as a neuromodulator of headsweep CPG networks. Intriguingly, previous computational work predicted the presence of network oscillators subject to neuromodulation as core circuit motifs controlling headsweeps (32,33). Our work provides experimental support for this conceptual framework. Recent modelling work shed light onto the roles inhibitory motifs and hemi-segmental oscillators play in generating headsweeps (34). However, to date, the exact cellular architecture of the headsweep CPG network remains largely unknown and the functional role of specific CNS regions, particularly the brain, are debated (20,35). Our imaging work identifies dopaminergic TH-neurons with activity patterns that encode dynamic features of headsweep circuit activity and may provide entry points into understanding the functional architecture of headsweep circuits. Breaking up the TH-GAL4 expression pattern to dissect the roles of specific cell types remains a question for future study. Functionally, dampening headsweep motifs may discourage exploratory headsweep behavior in favor of directed forward locomotion. That said, the lack of modulatory effects on headsweep behavior in intact animals suggests dopamine may constrain activity in a context-dependent manner. Removing sensory context with the isolated CNS preparation may bias the state of the headsweep CPG to be more receptive to dopaminergic modulation. Conversely, the added sensorimotor context of the environment may suppress dopamine’s dampening of headsweeps (i.e., exploratory behavior) in intact larvae. This highlights the value of using both isolated and intact preparations to understand the complete roles of neuromodulators.

Dopamine also modulated the generation of forward locomotion. Short-term global applications of high dopamine and L-dopa concentrations biased the isolated CNS towards increasing fictive forward locomotion. Increased fictive forward wave frequency was accompanied by decreased fictive backward waves in dopamine, supporting the hypothesis of suppressing competing motor programs via inhibitory motifs (34). These results share some similarities with adult *Drosophila* work showing that enhancing dopaminergic signaling promotes locomotion. In decapitated adult flies, application of 10mM dopamine and 5mM D2-like agonist promotes locomotion, while application of D2-like or D1-like antagonists reduces locomotion (36). Furthermore, RNAi-mediated reduction in D2-like receptor expression leads to reduced locomotion in adult flies (37) and mutant flies lacking the gene encoding tyrosine hydroxylase show reduced activity and locomotor deficits (38). By trialing a range of dopamine concentrations, we also found dose-dependent effects. Headsweep suppression and forward wave generation were stimulated under 250µM and 500µM dopamine but not 100µM. Thus, an activation threshold, or precise balance of D2-like to D1-like receptor activation may be required to produce these effects. Similar concentration dependency is observed in vertebrates, where lowered dopamine acts via inhibitory D2-like receptors to reduce locomotion, whilst high levels recruit excitatory D1-like receptors to increase locomotor episodes (39–41). In larval *Drosophila* research, dopaminergic signaling has previously inhibited locomotor activity (18,42) or was suggested to not be required for locomotion generation (19). It is important to note that these studies genetically manipulated dopamine signaling pathways and receptor expression throughout development. However, *Drosophila* have been shown to adapt to altered biogenic amine levels over time. For example, larvae with mutated *Drosophila* vesicular monoamine transporters have reduced motor neuron electrical activity, but compensatory increased motor neuron glutamate release (43). Moreover, the short and long-term effects of amines in other invertebrates depend on the administration method (44). Our acute manipulations of dopamine (minutes timescale) on the isolated CNS led to similarly short-term changes in motor neuron activity, providing a counterpoint to results from long-term manipulations. Therefore, we believe manipulation timescales have a fundamental effect on the actions of neuromodulators. Since locomotion requires modulation to adapt to rapid changes in environment, predation and food, faster timescale manipulations may uncover previously unseen, more ecologically relevant effects of modulators like dopamine.

Our behavioral results give further insight into the relationship between neuromodulation and the sensory environment, and suggest that dopamine does not promote/inhibit locomotion, but instead modulates CPG outputs based on current sensory and motor context. Optogenetic activation of TH-neurons during surface crawling increased forward wave duration and decreased wave frequency. Conversely, stimulation of the same neurons during tunneling increased wave frequency and thus total time spent tunneling. Strikingly, optogenetic inhibition of TH-neurons had no effect on surface crawling but impacted substrate tunneling. This highlights the importance of studying neuromodulation across multiple behavioral contexts to help avoid false negatives in optogenetic studies. Our results have interesting implications for interpreting optogenetic screens carried out within and outside *Drosophila* research. Overall, dopaminergic signaling appears to form an important part of the production of tunneling (but not crawling) locomotor behavior, with a role in coordinating a network shift toward enhancing tunneling. This is further evidenced by the presence of posterior asymmetries, a feature exceedingly rare during crawling and predominantly found in larval tunneling (unpublished observations), but readily induced in crawling larvae upon activation of TH-neurons. Higher posterior asymmetry frequency in these larvae vs normal tunneling larvae may reflect higher wave frequency in crawling over tunneling. Functionally, posterior asymmetries may play a stabilizing/anchoring role to help propel larvae into the substrate, supporting previous biomechanical studies (45) and similar adaptive functions in other invertebrates against challenging environments (46,47). To our knowledge, this is the first detailed quantification of locomotor parameters during larval tunneling in *Drosophila*.

State-dependency in neuromodulation is a well-evidenced concept (48–51), and dopamine is no exception. The outcome of dopamine modulation in the isolated mouse spinal cord depends on baseline network excitability (8). Indeed, an intricate interplay between internal state and neuromodulation often dictates behavioral choice (52–54) and behavioral transitions due to dopaminergic neuromodulation have been widely evidenced in invertebrates (9,10). Moreover, dopamine is involved in both producing and enhancing slow locomotion rhythms in invertebrates and vertebrates (55–57). Taken altogether, our results suggest dopamine may induce shifts from crawling to tunneling motor programs in larval *Drosophila.* More specifically, in larvae, we argue dopamine is not an initiator for tunneling behavior, but an initiator of tunneling rhythms. This is because TH-neuron activation does not trigger crawling larvae to tunnel, but instead slows overall rhythms and pushes wave frequencies closer to ∼0.2Hz, which is the tunneling frequency. Whereas in submerged larvae, the switch in forward locomotion strategy from crawling to tunneling due to the demands of a confined environment means, instead of slowing rhythms, dopamine augments rhythm frequency to enhance tunneling. The lack of change in wave duration upon activating TH-neurons during tunneling suggests there is an upper boundary for dopamine’s ability to increase wave duration. This is interesting as it suggests dopamine increases wave duration only far enough to bring it into an ecologically-useful pattern, rather than slowing wave duration in a dose-dependent manner. Dopamine’s activity may modulate neurons involved in the production of crawling to produce a tunneling gait, or it may activate an independent tunneling network, which competitively inhibits the crawling gait. Dopamine’s role in the production of tunneling provides an entry point for addressing the nature of gait-related neural architecture. Future studies could use dopamine receptor distribution to identify neurons of interest that may mediate motor pattern dynamics. Larval locomotion is likely produced by interactions between chains of coupled oscillators in each hemi-segment (20,34); however, the exact architecture of tunneling-related circuits remains unclear.

Our results also raise the question of whether forward locomotor activity in fictive preparations, which is considerably slower than *in vivo* crawling, represents crawling or tunneling. Our pharmacological experiments showed that dopamine and L-dopa increased fictive forward locomotion frequency, akin to how TH-neuron activation increased wave frequency during tunneling. At the neuromuscular level, recent work has shown that larval lateral transverse (LT) body wall muscles do not contract during unrestrained locomotion (58), which differs from results in animals constrained to linear channels (59) and isolated CNS imaging of motor neurons innervating LTs (60). Fictive locomotion is often used as a proxy for larval crawling (61,62). It is possible that this difference in neuromuscular recruitment might represent a change in gait. Fictive waves are also much slower (∼11s) than intact crawling (∼1s) waves (61), which is also a key difference between crawling and tunneling behaviors. Previous work has proposed that the differences seen between intact crawling and isolated fictive locomotion are caused by differences in excitatory sensory feedback in each context (63,64). Our work highlights the possibility that these differences may also be related to a shift from a crawling to tunneling gait.

Overall, this paper used fictive and intact preparations to illustrate how dopamine modulates the larval *Drosophila* motor network to enhance tunneling. Dopamine increased fictive forward locomotion and suppressed headsweep motor programs, potentially shifting the network toward direction-constrained locomotion, critical for effective tunneling. Behaviorally, dopamine activation indeed slows crawling rhythms on the surface, increases the frequency of tunneling-related phenotypes and increases wave frequency underground, suggesting dopamine mediates a coordinated network effort toward enhancing tunneling motor programs. Finally, we emphasize the importance of using acute manipulations, complementary preparations and different environmental/behavioral contexts to study adaptive locomotion, as neuromodulation itself is heavily modulated by state and sensorimotor context.

## Data Availability

Data and analysis code are available here: https://doi.org/10.17630/685c0adc-20d9-4ea6-ac86-cfa33e1bc661

## Supplemental Material

Supplemental Video. S1: https://youtu.be/lPgTfcqAfv0

Supplemental Video. S2: https://youtu.be/sjMBQGuaHDU

## Acknowledgements

The authors thank Dr Albert Cardona for hosting J.M. in his lab and teaching connectomics techniques, Cairn Research Ltd. for hosting J.M. for an internship and providing equipment, Dr Karen Hibbard for fly lines, Dr Pierce Mullen for coding advice, and current and past members of the Pulver Lab and Zwart Lab for their support.

## Author Contributions

J.M., S.R.P. and B.X.Y. conceived and designed research, J.M. and B.X.Y. performed experiments, J.M., B.X.Y., S.R.P. and M.F.Z. analyzed data, J.M., B.X.Y., M.F.Z. and S.R.P. interpreted results of experiments, B.X.Y. and J.M. prepared figures, B.X.Y. and J.M. drafted manuscript, B.X.Y., M.F.Z., S.R.P. and J.M. edited and revised manuscript.

